# The spatial structure of feedforward information in mouse primary visual cortex

**DOI:** 10.1101/2019.12.24.888156

**Authors:** Jun Zhuang, Rylan S. Larsen, Kevin T. Takasaki, Naveen D. Ouellette, Tanya L. Daigle, Bosiljka Tasic, Jack Waters, Hongkui Zeng, R. Clay Reid

## Abstract

Location-sensitive and motion-sensitive units are the two major functional types of feedforward projections from lateral genicular nucleus (LGN) to primary visual cortex (V1) in mouse. The distribution of these inputs in cortical depth remains under debate. By measuring the calcium activities of LGN axons in V1 of awake mice, we systematically mapped their functional and structural properties. Although both types distributed evenly across cortical depth, we found that they differ significantly across multiple modalities. Compared to the location-sensitive axons, which possessed confined spatial receptive fields, the motion-sensitive axons lacked spatial receptive fields, preferred lower temporal, higher spatial frequencies and had wider horizontal bouton spread. Furthermore, the motion-sensitive axons showed a strong depth-dependent motion direction bias while the location-sensitive axons showed a depth-independent OFF dominance. Overall, our results suggest a new model of receptive biases and laminar structure of thalamic inputs to V1.

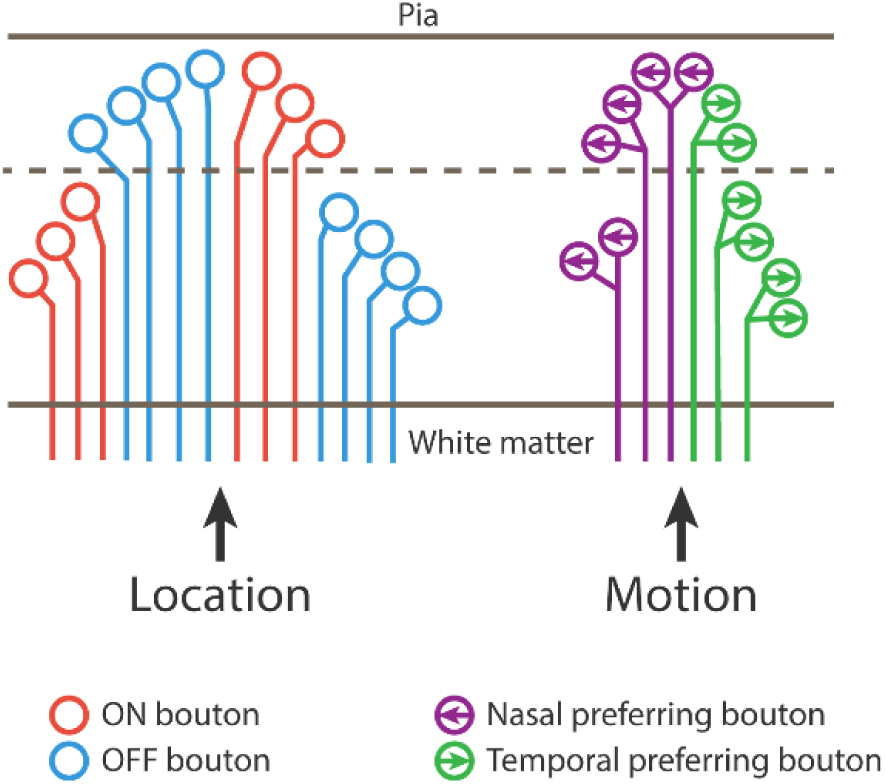

## Introduction

Parallel pathways along hierarchical processing stages are a prominent feature of early visual systems in carnivores (Stone et al., 1979; Lennie, 1980) and primates (Nassi and Callaway, 2009), where structured thalamocortical projections from dorsal lateral geniculate nucleus (LGN) to primary visual cortex (V1) serve as major inputs to cortical computations and shape cortical single-cell response properties (e.g., orientation selectivity, Hubel and Wiesel, 1962; Reid and Alonso, 1995) as well as the functional architecture at larger scale (Jin et al., 2011; Kremkow et al., 2016).

In recent years, the mouse visual system has become a popular model system to study visual physiology. When compared with higher mammals, the mouse LGN shows more diverse response properties. Particularly, besides cells with canonical ON and/or OFF concentric receptive fields (Grubb & Thompson, 2003; Denman et al., 2016, Suresh et al., 2016), a significant portion of the cells in mouse LGN are sensitive to motion (Marshel et al., 2012; Piscopo et al, 2013; Zhao et al., 2013; Durand et al., 2016) which may be inherited from its retinal inputs (Huberman et al, 2009; Cruz-Martin et al, 2014; Rompani et al., 2017; Dhande et al., 2015; Román Rosón et al., 2019). How this motion information is transmitted at the thalamocortical stage is less clear. It has been proposed that the motion information is conveyed by the direction-selective (DS) cells from LGN shell to the superficial layer of V1 while the location information is conveyed by cells in LGN core with concentric receptive fields to deeper layers (Cruz-Martin et al., 2014, Seabrook et al., 2017), forming parallel pathways similar to those in carnivores and primates. But the evidence for this model is scarce and controversial, while one study found mouse layer 4 in V1 receives slightly less DS inputs than layer 1 (Kondo et al., 2016), another study showed layer 4 receives the same amount of DS inputs as superficial layers (Sun et al., 2016).

To investigate the spatial distributions of LGN inputs in V1, we used genetic and viral tools to label mouse LGN neurons with calcium indicator and systematically measured the response properties of their axon terminals in V1 at different depths. We show that the motion-sensitive axons and location-sensitive axons represent two major functional types that differ significantly in functional properties, spatial distribution, and axonal morphology, demonstrating that there are at least two parallel thalamocortical pathways in mouse visual system.

## Results

### Selective labeling and functional imaging of LGN axons in V1

To image LGN axons in V1, we used the Vipr2-IRES2-Cre-neo transgenic mice. Crossing Vipr2-IRES2-Cre-neo mice with reporter line Ai14 drove tdTomato expression in the cells from LGN, ventral posterolateral nucleus, ventral posteromedial nucleus of thalamus, suprachiasmatic nucleus and central amygdala nucleus but not in the lateral posterior nucleus (LP) of thalamus and cortex (Figure 1A left, Allen Brain Atlas Transgenic Characterization Experiments: <u>576523754</u>, <u>576524006</u>, <u>577720920</u>, <u>577721229</u>). In V1, densely labeled axons were visible in layer 1 through layer 4 (Figure 1A right). Among the tdTomato positive nuclei, only LGN projects to V1, suggesting that the labeled axons in V1 were from LGN. To deliver calcium indicator for *in vivo* imaging, we injected a mixture of AAV viruses containing Cre-dependent GCaMP6s and Cre-independent mRuby3 directly into the LGN of Vipr2-IRES2-Cre-neo mice (Figure 1B). Three weeks after injection, the somatic expression of GCaMP was restricted in LGN while mRuby was expressed along the injection track through cortex and hippocampus (Figure 1C left). Importantly, the axons from these LGN cells were also labeled with GCaMP throughout superficial and middle layers in V1 (Figure 1C right).

**Figure 1.**
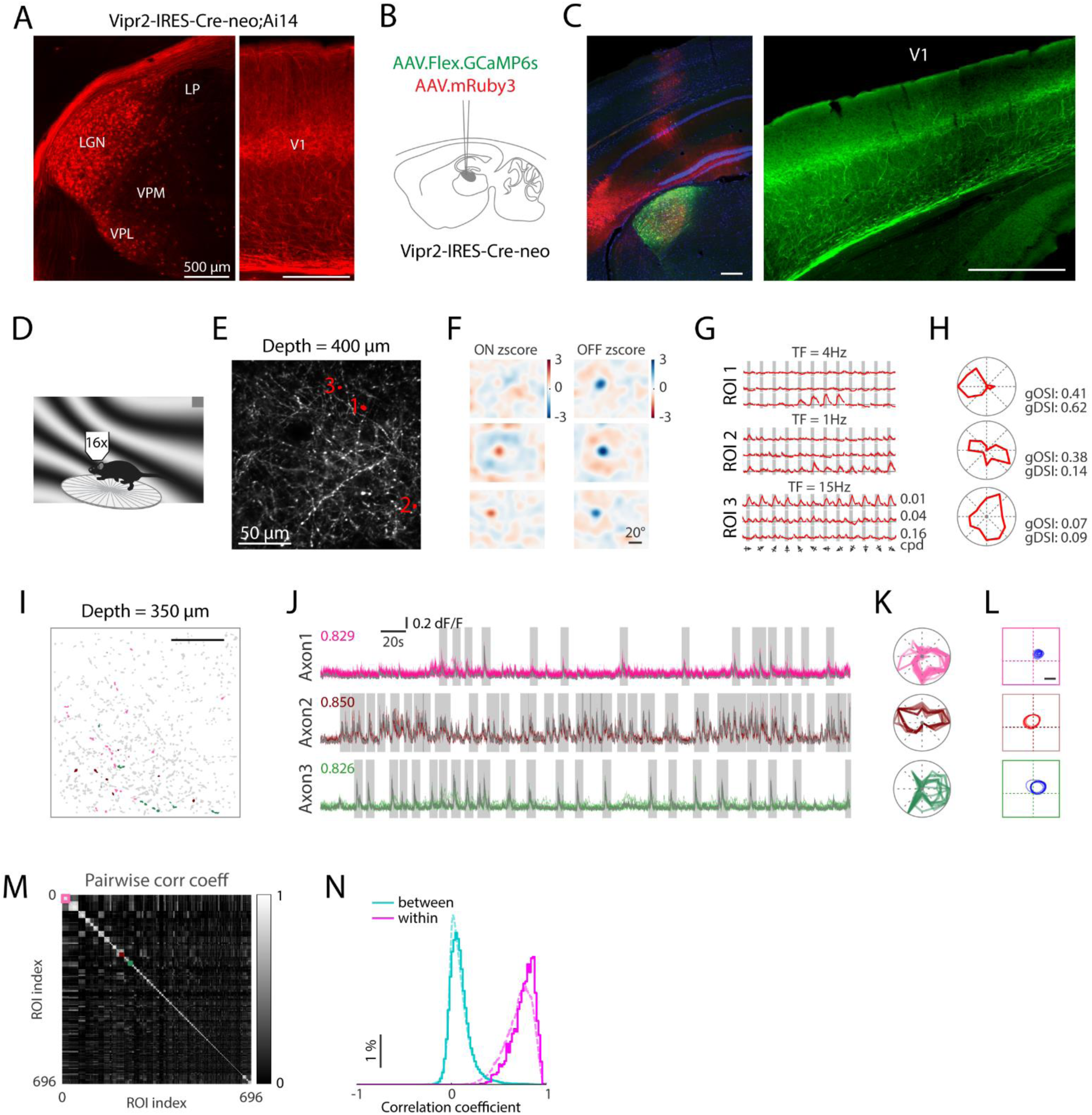
Labeling strategy, calcium imaging and bouton clustering for LGN axons in V1. (A) tdTomato positive cells in thalamus (left) and axons in V1 (right) from a Vipr2-IRES-Cre-neo;Ai14 mouse. (B) Labeling strategy for calcium imaging. (C) GCaMP (green) and mRuby (red) expressions in a mouse prepared as in (B). (D) Sketch of in vivo imaging setup. (E) Mean projection of calcium activities from an example imaging plane at 400 μm below pia. (F) The ON and OFF zscore receptive fields of the ROIs marked in (E). (G) The averaged calcium responses of ROIs marked in (E) to different grating conditions. Only the peak TFs for each ROI are shown. (H) The direction tuning curves of the ROIs marked in (E). (I) Another example imaging plane at 350 μm below pia showing all active ROIs (grey) and 3 example clusters after axon grouping (colored). (J) Calcium traces from the three marked axons. Colored traces: Traces from individual boutons. Gray traces: merged traces for the cluster. Vertical gray boxes: detected events. Numbers: mean pair-wise correlation coefficients. (K) Superimposed orientation/direction tuning curves of individual boutons from each cluster. (L) Superimposed receptive fields of individual boutons from each cluster. (M) Clustered correlation coefficient matrix for the example imaging plane in (I). Colored boxes: clusters marked in (I). (N) Normalized distribution of correlation coefficients for bouton pairs within clusters and between clusters. Solid lines: data from the imaging plane in (I). Dashed lines: data from all imaging planes.

We then performed two-photon calcium imaging of LGN axons 50-400 µm below the pia in awake mouse V1 (Figure 1D). The imaging was done in a columnar fashion: for a given cortical location, calcium signals from boutons were collected from 3 to 8 planes at different depths under pia. In total, 26 volumes (158 image planes from 6 mice) were imaged and calcium activity traces were extracted from 50171 regions of interest (ROIs) representing putative boutons (Table S1). During each imaging session, we measured spatial receptive fields (RFs), tuning for orientation/direction, spatial/temporal frequency (SF/TF), and spontaneous activity. Many ROIs showed strong calcium activities (Figure 1E and Video S1), significant spatial RFs (Figure 1F and S2A), and responses to gratings (Figure 1G-H). In some experiments, aberrations were reduced with adaptive optics, thereby increasing fluorescence intensity and signal-to-noise ratio. The response properties measured with and without adaptive optics were substantially similar (Figure S1), so they were pooled together. In summary, our labeling strategy enabled us to selectively label LGN neurons and characterize the functional properties of their axons in V1.

### LGN boutons can be grouped into axons based on highly correlated activities

To map individual sources of LGN inputs to V1, it is necessary to distinguish boutons that are from single axons from boutons that are from different axons. In each imaging plane, there were subgroups of ROIs showing highly correlated activities indicating that they were likely from same axons (Figure 1I-J, and Video S1). To identify these subgroups, we calculated an event-based pairwise correlation between all pairs of boutons, followed by a hierarchical clustering procedure (Methods, modified from Liang et al., 2018). Overall, the clustering faithfully preserved the pairwise correlation distance (Figure 1M) and the correlation coefficients from bouton pairs within and between clusters showed non-overlapping distribution (Figure 1N). The boutons from single clusters showed highly correlated calcium activities (Figure 1J) and near-identical response properties (Figure 1K-L), further confirming their high likelihood of being from single axons. We found the clusters generated with full calcium traces also represented the correlation structure of spontaneous activities (Figure S3), thus all imaging planes were processed using the full calcium traces. We use “bouton” when referring to individual ROIs, “axon” to indicate bouton cluster with high temporal correlations, and “unit” to refer to either entity. For each axon, we used spatially summed calcium traces from all its boutons to calculate its response properties (Method).

### Two populations of LGN axons with distinct functional and structural properties

We found no depth-dependent differences in population average of direction/orientation selectivity, preferred TF, and RF strength, while preferred SF was slightly higher at greater depths (Figure S4). These results were consistent with a previous report (Sun et al, 2016), but did not support the prediction of depth-dependent projection of motion-sensitive inputs (Seabrook et al., 2017). We then focused our analysis on motion sensitivity and location sensitivity, which are most likely to represent parallel pathways suggested by previous studies (Piscopo et al., 2013, Suresh et al., 2016). We found that location-sensitive boutons (boutons with significant RFs, RF boutons) were more likely to have low direction selectivity than the boutons without RFs (Figure 2A) while the motion-selective boutons (boutons with gDSI > 0.5, DS boutons) were more likely to have low RF strength than non-DS boutons (Figure 2B). Only a small portion of boutons were both location-selectivite and motion-selectivite (RFDS boutons 5.32% of all boutons, Figure 2C-D), indicating DS boutons and RF boutons are largely separate functional types. Consistent with Figure S4, we found no depth dependence in the number of DS and RF boutons up to 400 µm from the pia surface (Figure 2E). Similar functional dichotomy was also found in axons (Figure S5A-D).

**Figure 2.**
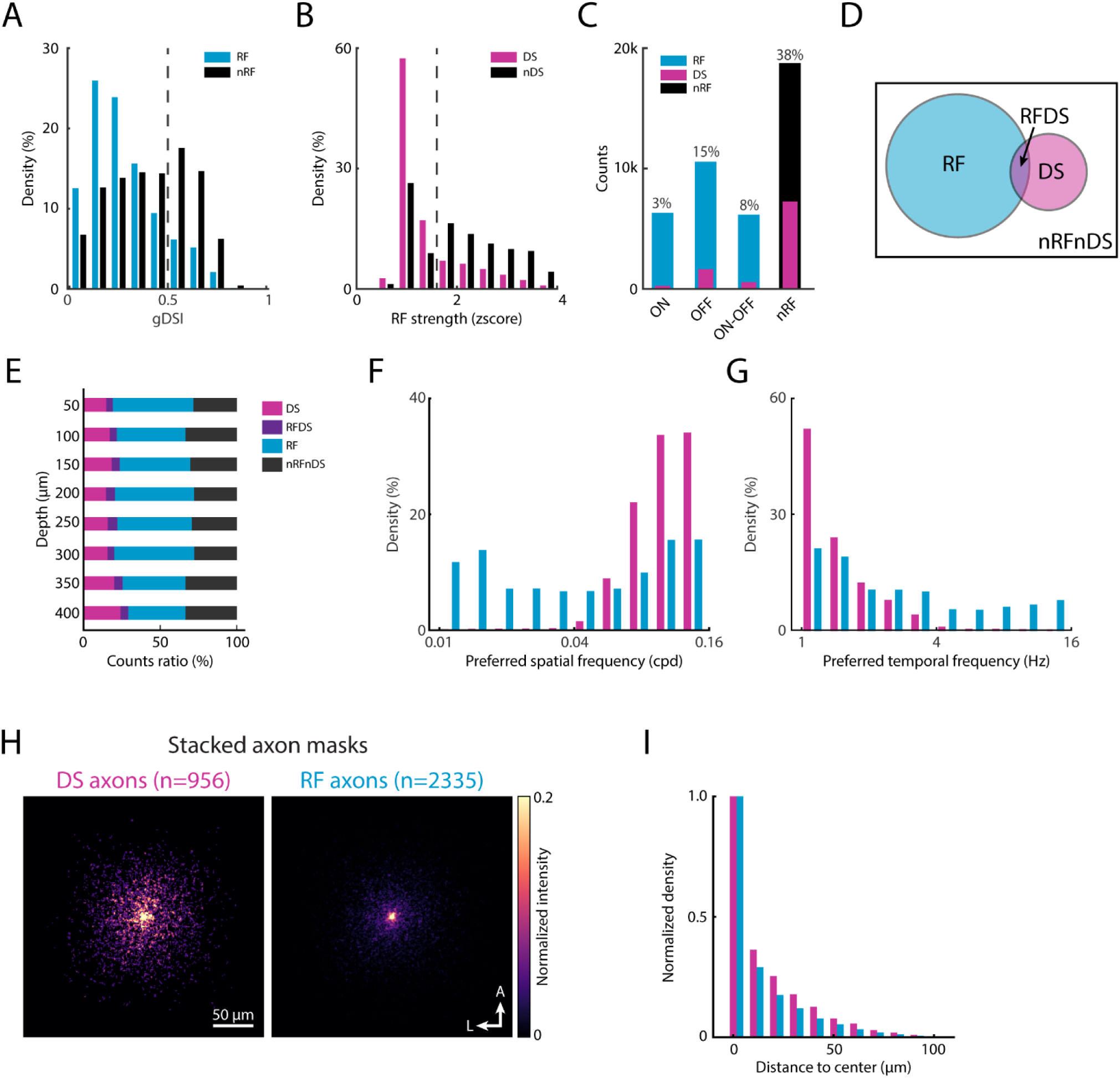
Motion-sensitive and location-sensitive axons are two major functional types of LGN inputs with distinct response properties and axon morphologies. (A) Distribution of gDSI of boutons with (RF) or without (nRF) significant spatial receptive fields. Dashed line: DS threshold. (B) Distribution of receptive field strength of DS and non-DS (nDS) boutons. Dashed line: RF threshold (C) Counts of each functional type. RF group was broken into ON, OFF and ON-OFF groups. Numbers: the portion of DS units in each group. (D) Venn diagram describing relations among the four non-overlapping groups. (E) The ratio of each of these four groups across depths. (F) and (G) Preferred SF and TF distributions, respectively. (H) Stacked axon masks for both DS axons and RF axons. (I) Normalized bouton distribution by distance to axon center.

Do DS and RF groups differ in other aspects? We first compared the SF and TF tuning between DS and RF boutons. We found that while the distribution of preferred TF and SF from RF boutons spread widely across the spectrum tested, DS boutons preferred only high SFs and low TFs (Figure 2F-G, preferred SF: DS vs. RF, 0.102 ± 0.036 vs. 0.004 ± 0.063 cpd, U=188907, p<10^-100^; preferred TF: DS vs. RF, 1.45 ± 0.59 vs. 2.56 ± 2.80 Hz, U=9421173, p<10^-100^, Mann Whitney U test). Similar results were found in axons (Figure S5E-F). Secondly, we compared the normalized population axon masks between these two groups and found that the DS axons had a greater bouton spread than the RF axons (Figure 2H-I, mean distance from bouton to axon center, DS axons: 42.2 ± 27.3 µm, n=4093; RF axons: 39.7 ± 27.9 µm, n=9423; U=18121755, p=1.22×10^−8^, Mann Whitney U test). Consistently, the DS axons also showed higher bouton counts, greater max bouton distances, and larger axon coverage area than RF axons (Figure S6). These results indicate that the DS axons might have more extensive axon arbors than RF axons.

In summary, our data showed there are at least two separate groups of LGN inputs with one carrying motion information (DS units), preferring high SF and low TF, and having wider axon arbors, and the other carrying location information (RF units), preferring a wider wide range of SFs and TFs, and having narrower axon arbors.

### Depth-dependent motion direction bias in DS group

We next examined the preferred direction distribution of DS boutons, which strongly biased towards nasal and temporal directions (Figure S7) consistent with a previous report (Marshel et al., 2012). Interestingly, in a given imaging plane, the DS population was often biased towards either nasal or temporal directions (e.g., the temporal direction in the example plane, Figure S2B). When broken into different depths, the direction bias showed a marked trend across cortical depth: it gradually switched from nasal direction at 50 µm to temporal direction at 400 µm (Figure 3A). To quantify this depth-dependent bias, we defined a metric: nasal-temporal index (NTI, Figure 3B inset). A negative NTI means more nasal-preferring units than temporal-preferring units in a given population and a positive NTI vice versa. Consistent with the observations above, the superficial (< 225 µm) boutons had a nasal bias (NTI = 0.245, Figure 3B magenta), and the deep (> 225 µm) boutons had a temporal bias (NTI = -0.259, Figure 3B, green). This trend was significant across all mice (Figure 3C, NTI: n=6, superficial vs. deep, 0.22 ± 0.09 vs. -0.11 ± 0.11, W=0, p=0.028) and all volumes (Figure 3D, NTI: n=17, superficial vs. deep, 0.25 ± 0.10 vs. -0.03 ± 0.12, W=17, p=0.005), and was insensitive to the DS threshold (Figure 3E). A similar depth-dependent bias was also found in DS axons (Figure S8).

**Figure 3.**
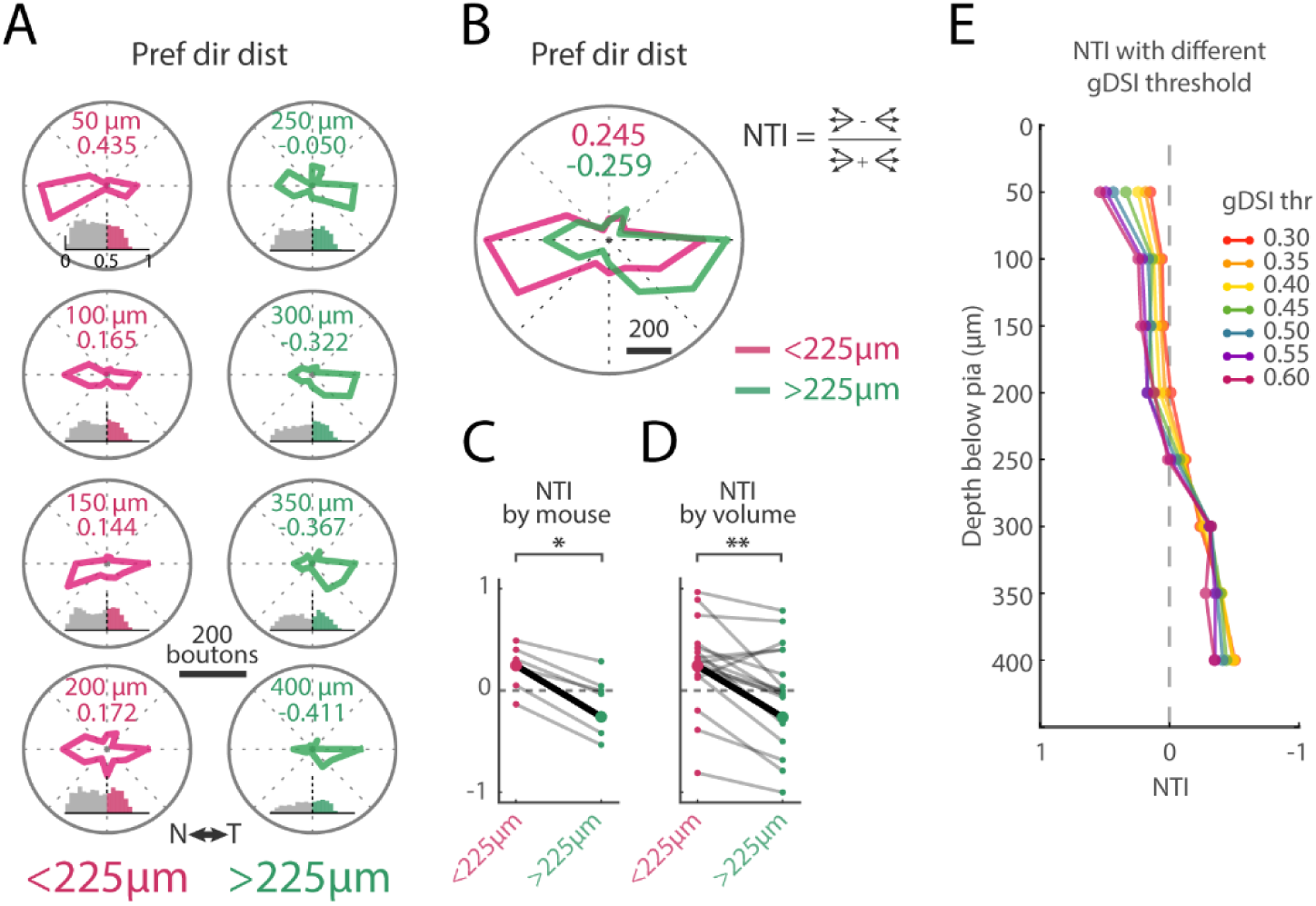
The motion-sensitive LGN boutons showed depth-dependent bias of motion direction preference. (A) Preferred direction distribution of DS boutons at each imaging depth. Numbers: NTI explained by the inset in (B). Histograms: distributions of gDSI at the given depth. Vertical scale: 100 boutons. Vertical dashed line: DS threshold. (B) Preferred direction distribution of superficial half vs. deep half of all DS boutons. (C) and (D), Superficial vs. deep NTI comparisons for each mouse and each volume, respectively. (E) NTI across depth with different DS thresholds.

### OFF dominance in RF group

Like previous reports in other species (Jin et al., 2008; Yeh et al., 2009; Jimenez et al., 2018), our RF boutons showed a strong OFF dominance. There were more OFF subfields than ON subfields per imaging plane (Figure 4A, all depths: n=121, ON vs. OFF, 54.5 ± 31.7 vs. 73.3 ± 52.1, W=1471, p=4.3×10^−8^) and this difference was consistent across cortical depth (ON vs. OFF, superficial: n=73, 53.6 ± 31.5 vs. 72.8 ± 52.7, T=536, p=1.3×10^−5^; deep: n=48, 55.8 ± 31.9 vs. 74.2 ± 51.1, W=234, p=8.1×10^−5^). Additionally, the OFF subfields were larger than ON subfields (Figure 4B, all depths, ON vs. OFF: n=121, 83.3 ± 2.0 vs. 102.1 ± 2.7 deg^2^, W=1480, p=1.1×10^−8^) also consistent across depth (ON vs. OFF, superficial: n=73, 81.5 ± 2.4 vs. 102.1 ± 3.8 deg^2^, T=532, p=6.8×10^−6^; deep: n=48, 86.0 ± 3.4 vs. 102.0 ± 3.5 deg^2^, W=240, p=3.6×10^−4^). A similar OFF dominance was also found in RF axons (Figure S9).

**Figure 4.**
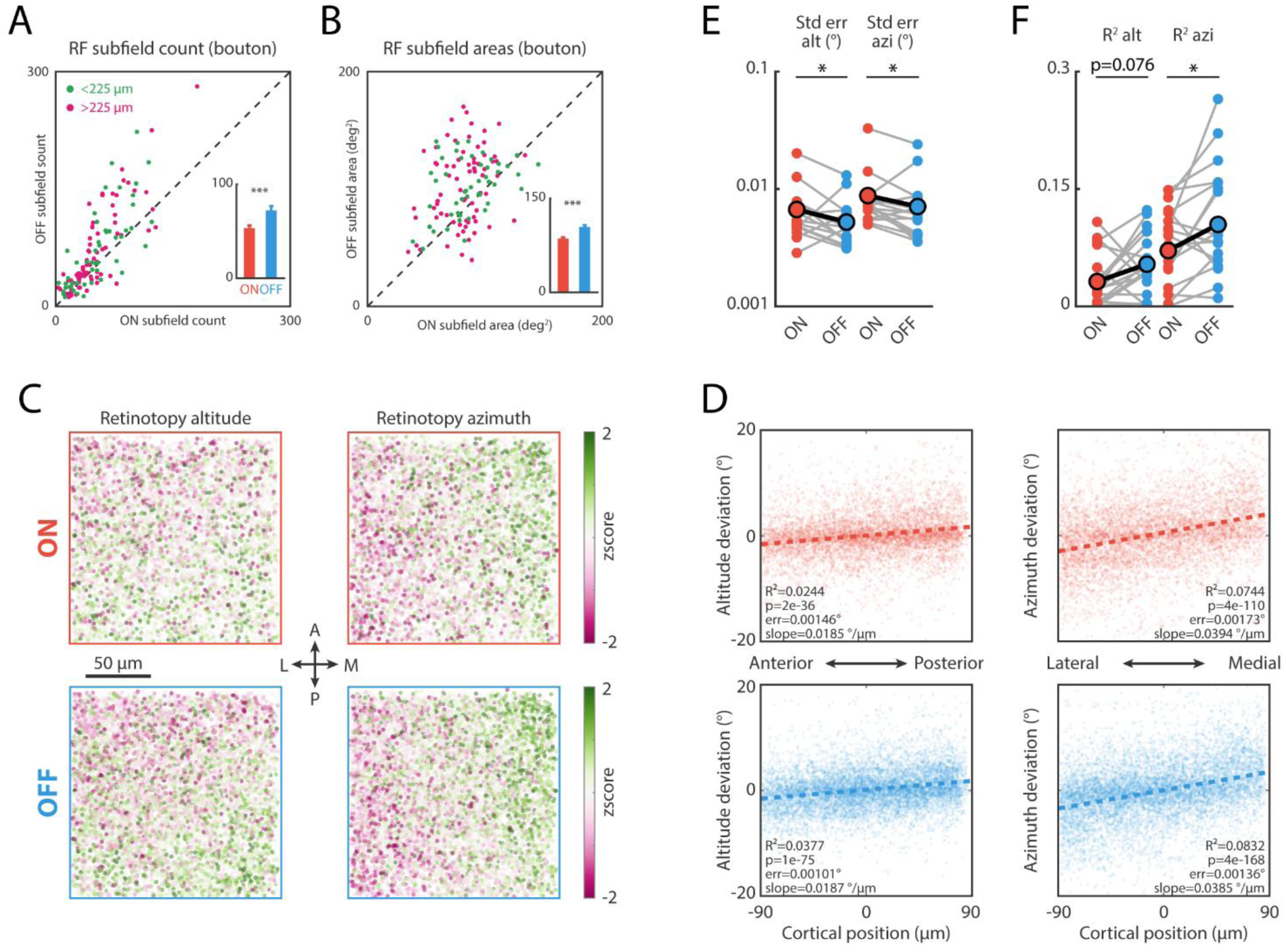
Location-sensitive LGN boutons were OFF dominant. (A) and (B) ON vs. OFF receptive subfield counts and areas of RF boutons, respectively. (C) Normalized retinotopy along cortical surface for RF boutons. (D) The linear regression of retinotopy location against the cortical distance. (E) and (F) ON vs. OFF regression standard error of estimate and R^2^ for each imaged volume, respectively.

Another form of “OFF dominance” that was prominent in other mammalian species (Kremkow et al., 2016; Lee et al., 2016), reduced retinotopic scatter for OFF cells, also presented in our data. We generated population retinotopic maps for both ON and OFF RF units by pooling maps from all imaging planes after alignment and normalization (Method). The pooled maps showed a clear retinotopic gradient (Figure 4C) consistent with the established retinotopic map of mouse V1 (Garrett et al., 2014; Zhuang et al, 2017), with altitude increasing along the anterior-posterior axis and azimuth increasing along the lateral-medial axis. More importantly, the retinotopic gradient appeared less noisy for OFF subfields than ON subfields. To quantify this difference, we performed linear regression of altitude against anterior-posterior locations and azimuth against lateral-medial locations. As expected, the regression fits better for the OFF retinotopy than for the ON retinotopy (Figure 4D). When the same analysis was performed on individual volumes, the differences between ON and OFF retinotopy were significant (except the R^2^ for altitude regression) and consistent with the entire population (Figure 4E-F, n=19, ON vs. OFF, standard error for altitude: 0.008 ± 0.001 vs. 0.006 ± 0.002 deg, W=37, p=0.02; standard error for azimuth: 0.011 ± 0.002 vs. 0.009 ± 0.004 deg, W=31, p=0.01; R^2^ for altitude: 0.032 ± 0.008 vs. 0.054 ± 0.029, W=51, p=0.076; R^2^ for azimuth: 0.071 ± 0.012 vs. 0.105 ± 0.049, W=35, p=0.016).

In summary, the OFF inputs in the RF group are more numerous, possess larger subfield, and have less retinotopic scatter than the ON inputs.

### Axon counts underlie functional specific bouton distribution

Finally, we asked what structure principle can give rise to the motion direction bias in DS boutons and the OFF dominance in RF boutons. One possibility is that axons with different functions might have different number of boutons (“bouton model”, Figure S10A and D, left). Alternatively, there could be different numbers of axons between different functional groups while having similar bouton number per axon (“axon model”, Figure S10A and D, right). The bouton model predicts different degrees of direction bias and OFF dominance for boutons and axons, while the axon model predicts similar degrees for boutons and axons. In favor of the axon model, our data showed the axons have similar NTIs (Figure S8 and S10B) and OFF dominance (Figure S9A and S10E) as the boutons. Our data also showed no difference in axon morphology between subtypes within DS and RF groups (Figure S10C and F), further supporting the axon model.

## Discussion

This study systematically investigated the functional properties and the spatial distributions of feedforward LGN axons in V1 with unprecedented labeling specificity. We showed the motion-sensitive (DS) axons and location-sensitive (RF) axons represent two major functional types that were largely distinct, with only ∼5% of units falling between groups (Figure 2C and S5C).

Compared to the RF axons, the DS axons preferred low temporal and high spatial frequencies and had a wider bouton spread. Furthermore, although neither type had depth-dependent distribution in V1, the DS group showed a depth-dependent bias in motion direction preference while the RF group showed a depth-independent OFF dominance. These findings support the notion that, in the thalamocortical projections of the mouse visual system, there are functionally and structurally distinct parallel pathways.

### LGN specific labeling

We found that Vipr2-IRES-Cre-neo mice allow superior specificity in LGN labeling than wildtype or Scnn1a-Cre mice used in previous studies (Kondo et al, 2016; Sun et al., 2016; Jaepel et al., 2017), because they effectively prevent off-target labeling in cortex and LP (Figure 1A and 1C), a region adjacent to LGN that projects substantially to V1 (Roth et al. 2016). This new labeling strategy may account for the different results between this study and previous studies, for instance if the Vipr2-IRES2-Cre-neo mice only label a subset of all LGN relay neurons. While we cannot disprove this from our data, we note that the proportions of the two main functional types in our dataset (RS axon: 53%, DS axon: 22%) are similar to what has been reported by previous electrophysiological studies (RF cells: 52%, DS/OS cells: 11% Piscopo et al., 2013; DS cells ∼25%, Zhao et al. 2013), which argues against significant functional bias in our labeling strategy.

### Parallel pathways

Based on recent understanding of parallel visual pathways in the mouse (reviewed in Seabrook et al., 2017), our RF units most likely correspond to X and Y units (Cleland et al., 1971; Hoffmann et al., 1972), and DS units correspond to W units in cat LGN (Wilson & Stone, 1975; Wilson et al., 1976). This is supported by our finding that DS axons had more extensive arbors than RF axons (Figure 2H-I and S6), consistent with the morphological differences between W and X/Y LGN axons in cats (Gilbert & Wiesel, 1979; Humphrey et al., 1985; Anderson et al., 2009). However, we found the DS units preferred low TF and high SF (Figure 2F-G), notably different from cat’s W LGN cells which prefer lower SF than X- and Y-cells (Sur & Sherman, 1982). Furthermore, we did not find that DS LGN cells project predominantly to superficial layers in V1 as predicted by the model (Seabrook et al., 2017), or mainly to layer 4 and 6 as has been seen for DS cells in other species (Hei et al., 2014; Bereshpolova et al., 2019). Instead, both RF and DS axons in our dataset distributed uniformly throughout all superficial depths measured (Figure 2E and S4), as had been reported before (Sun et al., 2016, cf. Kondo et al., 2016). Thus, the relationship between mouse LGN DS cells, W cells in the cat, and DS cells in other species needs further investigation.

Our statement that DS units tend to not have significant spatial RF should be taken with caution. The sparse-noise stimulus (Zhuang et al., 2013; Durand et la., 2016; de Vries et al., 2020) we used to map the spatial RF is effective if the unit responds to brief small flashes, but will perform poorly if the unit responds best to larger or more spatially structured stimuli, or requires a longer temporal duration. Nonetheless, our conclusions about parallel pathways are unlikely to be undermined because (1) a previous study with more structured mapping stimuli showed similar function dichotomy (Piscopo et al, 2013); and (2) we found other significant functional and structural differences between DS axons and RF axons (Figure 2F-I).

### Motion direction bias of DS axons

Consistent with previous studies (Marshel et al., 2012; Piscopo et al., 2013; Zhao et al., 2013; Durand et al., 2016; Sun et al., 2016; Kondo et al., 2016), we found that a significant population of LGN units are direction-selective (22% for axons, 23% for boutons, Figure 2C, Table S1). When pooling all our DS units together, the preferred directions strongly bias towards the nasal and temporal directions (Figure S7) consistent with Marshel et al. 2012 but unlike other studies (Sun et al., 2016: towards nasal direction; Kondo et al., 2016: towards four cardinal axes). These differences may be due to the different labeling strategies, visual stimuli, and analysis methods. The nasal/temporal direction biases have ethological relevance since they can be used to detect self-generated motion during active behavior, as has been demonstrated at both retina (Sabbah et al, 2017) and visual thalamus (Roth et al., 2016). Interestingly, our data show the direction bias changes systematically along with cortical depth (Figure 3 and S8). The functional relevance of this surprising organization is unknown and will be a topic for further investigation.

### OFF dominance

It is well established in the early visual system of carnivores and primates that the OFF responses are faster and stronger than ON responses (Jin et al, 2008, 2011; Yeh et al., 2009; Mazade et la., 2019; Kremkow et al., 2018). The OFF responses are also stronger in mouse V1 cells (Jimenez et al., 2018). We extend these findings to mouse LGN axons (Figure 4A-B and S9). Our data further show that the OFF subfields have less retinotopic scatter than ON subfields (Figure 4C-F), an organization underlying the structure of orientation maps in V1 of other species (Kremkow et al., 2016; Lee et al., 2016). Considering that mouse V1 lacks a systematic orientation map (Ohki et al., 2007), our results suggest such polarity-dependent retinotopy is conserved across various visual cortical architectures through evolution.

### Incomplete axon morphology

The axon morphology estimates of this study were performed on single two-photon imaging planes (∼ 190 x 190 µm in the XY plane and a few microns in Z). Given that the arbor of an LGN axon in V1 can be more than 500 µm in all three dimensions (Antonini et al., 1999), our measurements were substantially incomplete. We reasoned that, with such crude measurements, any subtle differences in axon morphology would fail to reach significance. However, to our surprise, the DS axons and RF axons showed significant differences consistently across all morphology measurements (Figure 2H-I and S6). This indicates that there should be dramatic differences between DS and RF axons in their full morphology, but further studies are needed to fully characterize the anatomical differences between the two physiological types.

## Methods

### Surgery and animal preparation

In total, six Vipr2-IRES2-Cre-neo mice (3 male, 3 female) were used in this study. The surgery included a stereotaxic viral injection and a cranial window/head-plate implantation. During the injection, a glass pipette back-loaded with a 1:3 mixture of AAV1-CAG-mRuby (custom made from plasmid addgene 107744, titer: 1.6×10^12^ vg/ml) and AAV1-Syn (or CAG)-FLEX-GCaMP6s (addgene: 100845-AAV1, titer 4.3×10^13^ vg/ml or 100842-AAV1, titer 1.8×10^13^ vg/ml, respectively) was slowly lowered into left LGN (2.3 mm posterior 2.3 mm lateral from bregma, 2.6 mm below pia) through a burr hole. 5 minutes after reaching the targeted location, 200 nL of virus mixture was injected into the brain over 10 minutes by a hydraulic nanoliter injection system (Nanoject III, Drummond). The pipette then stayed for an additional 10 minutes before it was slowly retracted out of the brain. Immediately after injection, a titanium head-plate and a 5 mm glass cranial window were implanted over left V1 (de Vries et al., 2020; for detailed protocol see Goldey et al., 2014) allowing *in vivo* two-photon imaging during head fixation.

After surgery, the animals were allowed to recover for at least 5 days before retinotopic mapping with intrinsic signal during anesthesia (for detailed retinotopic protocol, see Juavinett et al., 2017). After retinotopic mapping, animals were handled and habituated to the imaging rig for two additional weeks (de Vries et al., 2020) before *in vivo* two-photon imaging.

All experiments and procedures were approved by the Allen Institute Animal Care and Use Committee.

### Histology

To characterize the Cre expression pattern, Vipr2-IRES2-Cre-neo mice were crossed with the Ai14 reporter line (Madisen et al., 2010) to generate Vipr2-IRES2-Cre-neo/wt; Ai14/wt animals. Mice heterozygous for both Cre and the reporter were perfused and brains collected. Briefly, mice were anesthetized with 5% isoflurane and 10 ml of saline (0.9% NaCl) followed by 50 ml of freshly prepared 4% paraformaldehyde (PFA) was pumped intracardially at a flow rate of 9 ml/min. Brains were immediately dissected and post-fixed in 4% PFA at room temperature for 3-6 hours and then overnight at 4 °C. After fixation, brains were incubated in 10% and then 30% sucrose in PBS for 12-24 hours at 4 °C before being cut into 50 µm sections by a freezing-sliding microtome (Leica SM 2101R). Sections with LGN and V1 were then mounted on gelatin-coated slides and cover-slipped with mounting media (Prolong Diamond Antifade Mounting Media, P36965, ThermoFisher).

To verify the injection location and GCaMP expression, the brain tissues from all experimental mice were collected and sectioned with a similar procedure after all imaging sessions with additional steps with antibody staining to enhance the GCaMP signal before mounting and coverslipping. During antibody staining, sections containing LGN and V1 were blocked with 5% normal donkey serum and 0.2% Triton X-100 in PBS for one hour, incubated in an anti-GFP primary antibody (1:5000 diluted in the blocking solution, Abcam, Ab13970) for 48-72 hours at 4 °C, washed the following day in 0.2% Triton X-100 in PBS and incubated in a Alexa-488 conjugated secondary antibody (1:500, 703-545-155, Jackson ImmunoResearch) and DAPI. The sections were then imaged with Zeiss AxioImager M2 widefield microscope with a 10x/0.3 NA objective. Fluorescence from antibody enhanced GCaMP and mRuby3 were extracted from filter sets Semrock GFP-1828A (excitation 482/18 nm, emission 520/28 nm, dichroic cutoff 495 nm) and Zeiss # 20 (excitation 546/12 nm, emission 608/32 nm, dichroic cutoff 560 nm), respectively.

### *In vivo* two-photon imaging

Following the completion of surgical recovery, retinotopic mapping, and habituation (usually more than three weeks after initial surgery), thalamocortical axons from LGN labeled with GCaMP6s were evident in superficial- and mid-layers (0 – 500 µm) in V1 through the cranial window (Figure 1E). We choose 8 depths for in vivo two-photon imaging (50, 100, 150, 200, 250, 300, 350 and 400 µm below pia). Imaging was done in a columnar fashion: at each cortical location calcium activities at 3-8 depths were imaged plane-by-plane over multiple sessions with a conventional 2-photon microscope or in a single session with a multi-plane 2-photon microscope (described below). In both microscopes, a 16x/0.8 NA water immersion objective (Nikon 16XLWD-PF) was rotated to 24 degrees from horizontal to image visual cortex using a commercial rotating head (Sutter MOM). mitted light was first split by a 735 nm dichroic mirror (FF735-DiO1, Semrock). The short-wavelength light was filtered by a 750 nm short-pass filter (FESH0750, Thorlabs) and a 470-588 nm bandpass emission filter (FF01-514/44-25, Semrock) before collected as GCaMP signal, while the long-wavelength light was filtered by a 538-722 nm band-pass emission filter (FF01-630/92-30, Semrock) before collected as mRuby signal. Image acquisition was controlled using Vidrio ScanImage software for both scopes (Pologruto et al., 2003, Vidrio LLC). To maintain constant immersion of the objective, we used gel immersion (Genteal Gel, Alcon).

#### Single-plane two-photon imaging

Four mice were imaged with two-photon excitation generated by laser illumination from a Ti:sapphire laser (Coherent Chameleon Ultra II) tuned to 920 nm. A single z-plane (179.2 x 179.2 µm with 512 x 512 pixels resolution) was imaged for each session at a frame rate of about 30 Hz with an 8 KHz resonate scanner (Cambridge Technology, CRS 8K).To maintain constant imaging depth automatic z-drift correction functions were implemented for experiments using the MOM motors. Briefly, a correction z-stack (± 50 µm from targeted depth, 2 µm step depth) was recorded before each imaging session and, during the session, the current imaging plane was continuously compared to each plane in the correction z-stack. If a drift in depth was detected, the stage was automatically adjusted to compensate for the drift, thus maintaining constant imaging depth. We found this procedure crucial to our experiments since the boutons are small objects and a few-micron-drift in depth would result in imaging a different set of boutons.

#### Multi-plane two-photon imaging

The other two mice were imaged by a custom-built, multi-plane two-photon microscope (DeepScope, Liu et al., 2018) in which a liquid crystal spatial light modulator (SLM; HSP-512, Meadowlark Optics) shapes the pupil wavefront to implement fast-focusing and adaptive optics. The objective pupil was slightly underfilled (∼0.65 effective NA) and correction of systemic aberrations was performed with fluorescent beads to maintain near diffraction-limited focusing over a 200 um range. Two-photon excitation was produced by laser light from a commercial solid-state laser (Spectra-Physic Insight X3 laser) tuned to 940 nm. With this microscope, we simultaneously recorded calcium activity from planes at 5 different depths (50, 100, 150, 200, 250 µm) in single imaging sessions. Individual frames (125 x 125 µm with 512 x 512 pixels resolution) were acquired at an overall framerate of ∼37 Hz with a volume rate of 7.4 Hz. The DeepScope showed nearly zero z-drift for a prolonged duration (< 2 µm over 24 hours), so we did not implement z-correction in sessions using DeepScope.

In another set of experiments, we used DeepScope to assess the effect of adaptive optics adjusted on individual animals. The correction procedure was similar to the method described in Sun et al., 2016. For two mice, 1 µm beads (Thermo Fisher, F8821) were deposited on top of the brain surface under the coverglass during the initial surgery. Prior to the imaging session, modal optimization over 12 Zernike modes (up to j = 15 Noll ordering, excluding piston and tilt) was run to identify the SLM pattern that maximized the beads’ fluorescent signal. Then, during the imaging session, this SLM pattern was put on and off alternatively for consecutive two-photon imaging frames (one frame on, one frame off, repeated) and drifting gratings were displayed. After imaging, the interleaving movie was separated into two movies: one with adaptive optics and the other without. Bouton’s tuning properties were then extracted from each movie and compared against each other.

All imaging sessions were performed during head fixation with the standard Allen Institue Brain Observatory *in vivo* imaging stage (de Vries et al., 2020).

### Visual stimulation

All visual stimuli were generated and displayed by Retinotopic_Mapping python package (https://github.com/zhuangjun1981/retinotopic_mapping, Zhuang et al., 2017) over PsychoPy software (https://www.psychopy.org, Peirce et al., 2019) on a 24-inch LCD monitor (ASUS PA248Q, frame rate 60 Hz, 1920 x 1200 pixels, mean luminance 45.3 cd/m^2^) placed 15 cm from the mouse’s right eye (covering 120° x 95° of monocular visual space). We displayed locally sparse noise and/or full-field drifting grating in each imaging session to measure receptive fields and orientation/direction/spatial and temporal frequency tuning properties, respectively. In most sessions, we also displayed a five-minute full-field mid-luminance gray to measure spontaneous activity. For locally sparse noise, bright and dark squares (5° x 5°) were displayed in a random sequence on a grid tiling the entire monitor. At any given time, multiple squares could be displayed but the minimum distance between those squares should be no less than 50°. Each square lasted 100 ms and in total was displayed 50 times. For drifting gratings, the combinations of 12 directions (every 30°), 3 spatial frequencies (SF, 0.01, 0.04 and 0.16 cpd) and 3 temporal frequencies (TF, 1, 4, 15 Hz) were displayed. Each display lasted 1 second and was spaced by 1-second mean luminance grey period. In total, 3 x 3 x 12 + 1 (blank) = 109 conditions were randomly displayed in each iteration and the whole sequence contained 13 iterations. All stimuli were spherically corrected so that they were presented with accurate visual angles on the flat screen (Zhuang et al., 2017).

### 2-photon Image preprocessing

The recorded two-photon movies for each imaging plane were first temporally averaged across 5 frames (Sutter scope) or 2 frames (DeepScope), then were motion-corrected using rigid body transform based on phase correlation by a custom-written python package (https://github.com/zhuangjun1981/stia/tree/master/stia, Zhuang et al., 2017). We performed motion correction on the red channel (mRuby) and applied the offsets to the green channel (GCaMP). To generate regions of interest (ROIs) for boutons, the motion-corrected movies were further temporally downsampled by a factor of 3 and then processed with constrained non-negative matrix factorization (CNMF, Pnevmatikakis et al., 2016) implemented in the CaImAn python library (https://github.com/flatironinstitute/CaImAn, Giovannucci et al., 2018). The resulting ROI masks were normalized by subtracting the mean and dividing the standard deviation. Then pixels below 3 were set to 0. These ROIs were further filtered by their size (ROIs smaller than 1.225 µm^2^ or larger than 12.25 µm^2^ were excluded), position (ROIs within the motion artifacts were excluded) and overlap (for ROIs with more than 20% overlap, the smaller ones were excluded). For each retained ROI, a neuropil ROI was created as the region between two contours by dilating the ROI’s outer border by 1 and 8 pixels excluding the pixels within the union of all ROIs. The calcium trace for each ROI was calculated by the mean of pixel-wise product between the ROI and each frame of the movie, and its neuropil trace was calculated in the same way using its neuropil ROI. To remove the neuropil contamination, the neuropil contribution of each ROI’s calcium trace was estimated by a linear model and optimized by gradient descendent regression with a smoothness regularization (Zhuang et al., 2017; de Vries et al., 2020). Nonetheless, the estimates of neuropil contamination in our datasets were generally low (percentage of neuropil signal that contributed to the ROIs signal: 11.5 ± 0.5%) likely due to the small volume and sparse distribution of afferent axons in V1. As reported previously, a high skewness is an indication of active calcium activity (Mukamel et al., 2009, Dipoppa et al, 2018). The ROIs with skewness greater than 0.6 were defined as “active” boutons and were included in the following study, while others were excluded (Table S1). For each imaging session, a comprehensive file in Neurodata Without Borders (nwb) 1.0 format was generated to store and share metadata, visual stimuli, all preprocessing results, and final calcium traces using the “ainwb” package (https://github.com/AllenInstitute/nwb-api/tree/master/ainwb).

For the neuropil subtracted calcium traces with a minimum value below 1, an offset was added to the whole trace so that the trace minimum was 1. This procedure ensured a positive baseline for all units in the df/f calculation (below).

### Bouton clustering

The correlation-based bouton clustering procedure was based on previously reported algorithms (Liang et al., 2018) and the same procedure was performed on each imaging plane. First, we detected calcium events for each active bouton in the imaging plane. The calcium trace was smoothed by a gaussian filter with sigma of 0.1 seconds. Then event onsets were detected as up-crosses over a threshold of 3 standard deviations above the mean. A period 3 seconds before and after each onset was defined as an event window and the union of all event windows for a particular ROI was saved. Second, for any given pair of boutons, the union of event windows from both boutons was calculated and the calcium trace within the union window was extracted and concatenated for each bouton. Then a Pearson correlation coefficient of the two concatenated traces was calculated for this pair. By performing this procedure on every pair of active ROIs, we generated a correlation coefficient matrix for each imaging plane. We found the event detection important because it confined the correlation to the period only at least one bouton in the pair was active thus avoided correlating the noise during the inactive period. Third, the correlation coefficient matrices were further thresholded to reduce noise: the correlation coefficients for a given bouton were maintained if the coefficients were larger than 0.5 or if they exceeded 3 standard deviations above the mean value of all the coefficients between this bouton and all others. Otherwise, they will be set to 0. Third, a hierarchy clustering was performed to a given imaging plane using “1 – thresholded correlation coefficient matrix” as the distance matrix using Scipy.cluster.hierarchy library with a “weighted” method (https://docs.scipy.org/doc/scipy/reference/generated/scipy.cluster.hierarchy.linkage.html).

For each imaging plane, we calculated a cophenetic correlation distance to evaluate the clustering performance. Fourth, we use a threshold of 1.3 to separate clusters since ∼ 1.5 shows up as a relatively natural cut-off in the dendrograms. This threshold appeared somewhat conservative on the clustered correlation coefficient matrix (Figure 1M). Fifth, after clustering, each cluster was identified as a separate axon and its calcium trace was calculated as mean calcium trace of all boutons belonging to this cluster, weighted by the sum of their ROI weights.

For each detected axon, three values were extracted to estimate the morphology: (1) bouton number of this axon; (2) maximum distance among all bouton pairs belonging to this axon; (3) area of the convex polygon encapsulated by all the boutons belonging to this axon. For axons with only one bouton, the metrics 2 and 3 were set to be “nan” and were excluded from statistical analysis.

To generate stacked axon mask in Figure 2H, we first selected axons with at least two boutons for each group (DS axons or RF axons). Then the binary mask of each individual axon was rotated and scaled into the same standard orientation (up: anterior, left: lateral) and centered to its own center of mass. All centered axon masks were summed together to generate a summed population axon mask for each group. Each summed population axon mask was then divided by its own maximum value, generating a normalized population axon mask (Figure 2H). Axons with both significant RF and direction selectivity were excluded from this analysis to avoid double counting.

### Receptive field analysis

We calculated the units’ spatial receptive fields from their responses to the locally sparse noise stimulus. For a given square flash at a particular location with a particular sign, the calcium traces within a temporal window around the square onsets were extracted and aligned by the square onsets. An event-triggered average trace was then calculated as the mean trace across all aligned traces. The mean value of the averaged trace in the window [-0.5, 0] seconds from the onset was calculated as the baseline and event-triggered df/f trace was calculated as (average trace - baseline) / baseline. The response amplitude to this particular square was then calculated as the mean df/f in the window [0, 0.5] second from the stimulus onset. By repeating this procedure for all squares presented in locally sparse noise stimuli, a 2d amplitude map was generated tiling the entire visual space covered by the monitor for each sign (“ON” from bright squares and “OFF” from dark squares). From the amplitude map, a z-score map was calculated by subtracting the mean and dividing the standard deviation of the entire map. The z-score map was then smoothed by a Gaussian filter with a sigma of 1 pixel and upsampled by a ratio of 10 with cubic interpolation. The resulting map was defined as the receptive field map with a resolution of 0.5 degrees. The peak value of ON and OFF receptive field map combined was defined as RF strength plotted in Figure 2B and S4E. We defined a receptive field map with a peak value exceeding 1.6 as having a significant spatial RF. For each significant spatial RF, an RF mask was generated by thresholding either with a value of 1.6 (for maps with a peak less than 4) or with a value of 40% of its peak (for maps with a peak greater than 4, Zhuang et al., 2013). The RF area (plotted in Figure 4B and S9B) was calculated as the area of the binary mask after thresholding. The RF center was calculated as the location of the center of mass from the weighted mask after thresholding.

For the retinotopy analysis in Figure 4C-D, we rotated and scaled each imaging plane so that they had the same orientation of cortical surface (up: anterior, down: posterior, left: lateral, right: medial) with same pixel size. For each of these planes, we obtained all significant RFs from boutons in this plane and calculated their centers. The population RF was then calculated by summing all significant RFs and thresholding with the same steps described above. Then the relative RF location of each individual RF was calculated as the vector pointing from the population RF center to the individual RF center. The relative RF locations were then normalized by subtracting mean and dividing by standard deviation (z-score relative RF location). To generate the normalized retinotopy map for each imaging plane, the individual z-score relative RF locations were color-coded and plotted to the center of their boutons within the rotated and scaled imaging plane. Since the retinotopy gradient is relatively uniform across mouse V1 (Zhuang et al., 2017), these procedures allowed us to superimpose the normalized retinotopy maps from all imaging planes together. We performed this for ON and OFF subfields and altitude and azimuth separately generating four retinotopic maps with respect to each sign and visual space dimension (Figure 4C). To quantify retinotopic gradient and scatter, we performed linear regression between cortical locations and z-score relative RF locations from these four superimposed retinotopic maps with relevant axis (anterior-posterior axis against altitude and medial-lateral axis against azimuth, Figure 4D).

### Grating response analysis

The units’ responses to drifting gratings were analyzed by the event-triggered average procedure similar to the RF response analysis. The difference is the temporal window to calculate response was [0, 1] second from the grating onsets since the duration of each grating was 1 second. We first define a unit having significant responses to the drifting gratings if it reached two criteria: 1) it responded to at least one grating condition with a z-score greater than 3 and 2) the Oneway ANOVA analysis of df/f responses across all grating conditions (including blank trials) had a p-value less than 0.01. For a unit that had a significant response to gratings, its peak condition was defined as the conditions with the highest df/f response. Its direction tuning curve was extracted as its df/f responses to different directions with the TF and SF the same as the peak condition and, if the minimum response from this curve was below zero, an offset was added to the whole curve so that the minimum was zero. Its SF and TF tuning curves were extracted by a similar procedure. From direction tuning curve, we calculated the global orientation index (gOSI) as

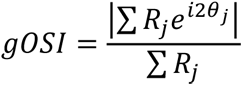

Where j represents different direction conditions, R represents df/f response in a given direction and θ represents the direction. The preferred orientation as the angle of 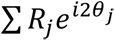. We calculated global direction index (gDSI) as

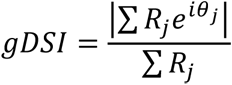

And the preferred direction as the angle of 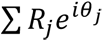. From the SF tuning curve, we calculated the preferred SF as

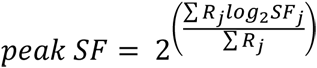

Where j represents different SF conditions. From TF tuning curve, we calculated the preferred TF as

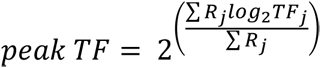

Where j represents different TF conditions. For the units that did not have significant grating response, all these measurements were set to be “nan”. We defined a unit to be direction-selective if it had a gDSI greater than 0.5 and orientation-selective if it had a gOSI greater than 0.3.

For each imaging plane, we calculated a nasal-to-temporal index (NTI, see main text):

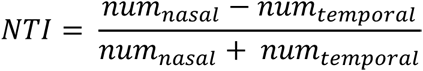

Where num_nasal_ is the count of DS units that had preferred direction in the range of [-45°, 45°] and num_temporal_ is the count of DS units that had preferred direction in the range of [135°, 225°]. Only the planes with num_nasal_ + num_temporal_ greater than 10 were included in the analysis. We also calculated an ON-OFF index:

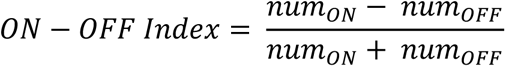

Where num_ON_ is the count of RF units that had only ON subfield and num_OFF_ is the count of RF units that had only OFF subfield. Only the planes with num_ON_ + num_OFF_ greater than 10 were included in the analysis.

The imaging preprocessing, nwb packaging, bouton clustering, receptive field analysis, grating response analysis, and functional type classification were performed by a custom-written python package “corticalmapping” (https://github.com/zhuangjun1981/corticalmapping/tree/master/corticalmapping).

### Statistics

To control the variability across imaging planes (different axon/bouton density, vasculature pattern, expression level), almost all statistics were extracted from each plane and separated for different functional types if necessary. For statistics that only one number can be drawn from each imaging plane (bouton/axon count, NTI, etc), they were presented as mean ± standard deviation. For statistics that a population distribution can be drawn from each imaging plane (gDSI, gOSI, peak SF/TF, RF strength, RF area, bouton per axon, max bouton distance, axon coverage area), mean for each plane was calculated first and mean ± standard error was reported across imaging planes. The comparisons between different functional types within imaging plane were performed by Wilcoxon rank-sum test, and comparisons between imaging planes were performed by Mann-Whitney U test since the distribution of the statistics were highly skewed and non-gaussian. We have verified our results of these two tests by paired t-test and independent t-test respectively, and they all agreed with the non-parametric tests (not shown). In the main text, the significant tests were Wilcoxon rank-sum test if not specified. All significance tests were two-tailed.

## Acknowledgments

Funding was provided by the Allen Institute for Brain Science and award number NS104949 from the National Institute of Neurological Disorders and Stroke. The contents of this manuscript are solely the responsibility of the authors and do not necessarily represent the official views of the National Institutes of Health and National Institute of Neurological Disorders and Stroke. We thank Dr. Roger Tsien for providing tdTomato construct. We thank many staff members of the Allen Institute, especially the In Vivo Sciences team for surgeries and the Manufacturing and Processing Engineering team for hardware support. We wish to thank the Allen Institute for Brain Science founder, Paul G. Allen, for his vision, encouragement, and support.

