## Supplemental Figures for "The spatial structure of feedforward information in mouse primary visual cortex"

| Depth (µm) | Number of planes | Number of active units | Number of units with significant RF | Number of units responsive to gratings | Number of DS units |
|---|---|---|---|---|---|
| 50 | 20 | 8120 (4002) | 2374 (1009) | 2396 (1237) | 716 (353) |
| 100 | 25 | 7347 (3862) | 1368 (620) | 1666 (887) | 576 (283) |
| 150 | 20 | 6924 (3330) | 1505 (662) | 1800 (921) | 629 (309) |
| 200 | 25 | 8469 (4128) | 2050 (917) | 2130 (1084) | 761 (334) |
| 250 | 20 | 7388 (3518) | 1828 (804) | 2030 (995) | 758 (319) |
| 300 | 20 | 6336 (3210) | 1707 (832) | 1536 (830) | 609 (284) |
| 350 | 9 | 3323 (1473) | 904 (407) | 1431 (637) | 568 (221) |
| 400 | 7 | 2264 (975) | 569 (250) | 982 (445) | 442 (171) |

Table S1. Total unit counts at different imaging depths. Unit counts are presented as "bouton number (axon number)".

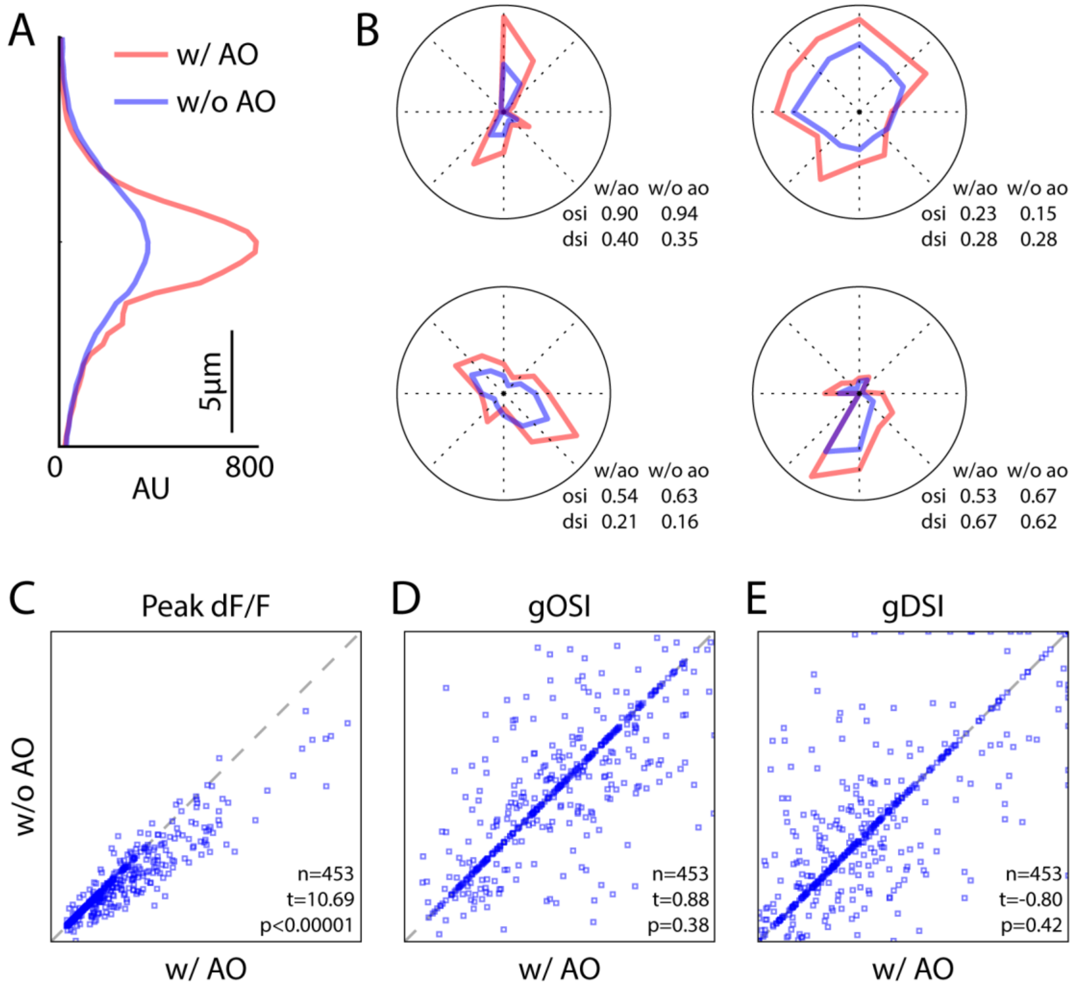


Figure S1. Comparisons of response amplitude, direction selectivity and orientation selectivity between conditions with and without adaptive optics (AO).

(A) Point spread function with and without AO by imaging 2 µm beads deposited between coverglass and brain surface in an awake mouse.

(B) Example orientation/direction tuning curves from 4 example boutons.

(C) - (E) Comparisons of peak dF/F, gOSI and gDSI of all recorded boutons. Two imaging planes in two mice (300 µm and 350 µm below pia). In AO conditions, laser beam wave front was corrected separately for each mouse. Significance test: paired t-test.

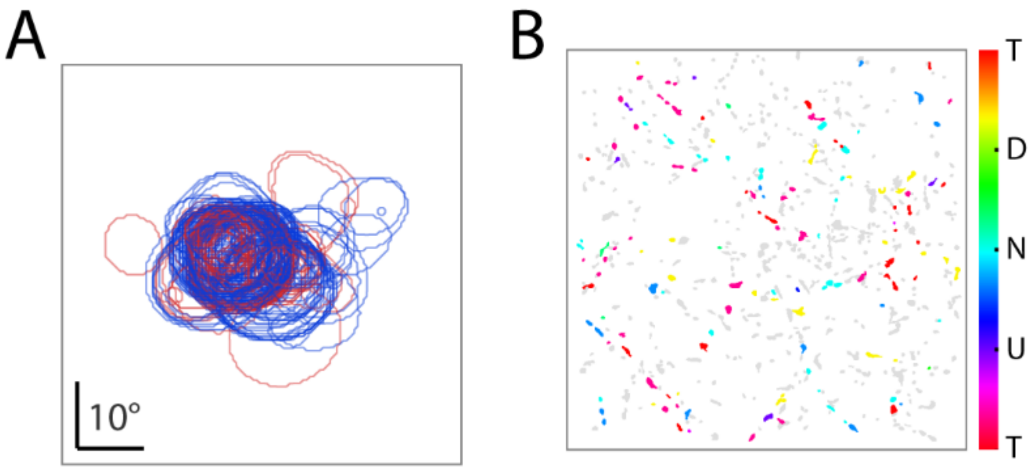


Figure S2. Receptive fields and preferred directions of all ROIs in the example imaging plane of Figure 1E

(A) All significant receptive fields. Each contour represents one subfield.

(B) Spatial distribution of preferred directions. Grey indicates no significant response to gratings or low direction selectivity. N, nasal; T, temporal; U, up; D, down.

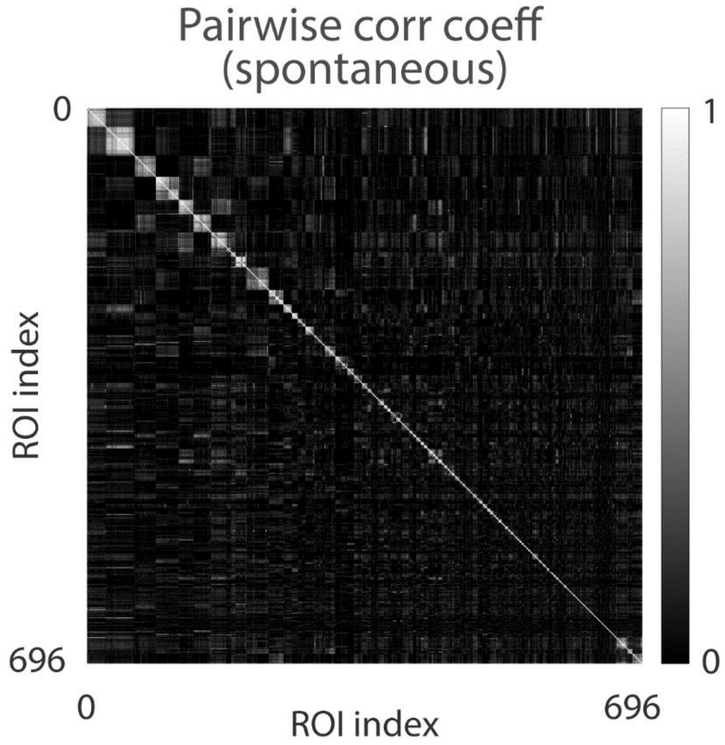


Figure S3. Correlation coefficient matrix calculated from only spontaneous activity sorted by the clusters generated in Figure 1M.

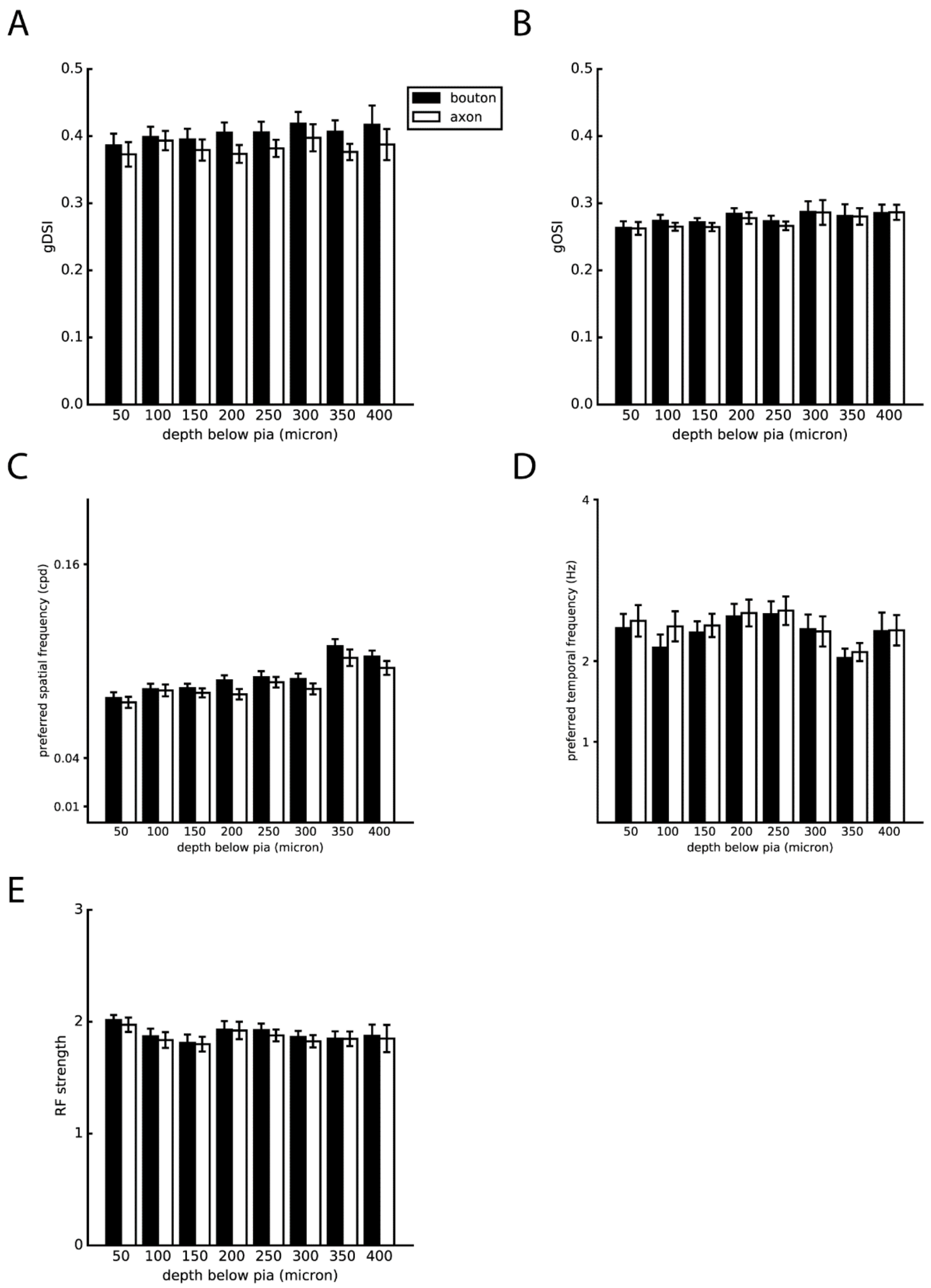


Figure S4. Response properties at different depth. For panel (A)-(D), only units with significant response to drifting gratings were included. For each depth, averages of a given response property per plane were calculated and the mean and s.e.m. of these averaged values are plotted here.

Statistics: global direction selectivity index, gDSI, boutons: 0.40 ± 0.003, F=0.34, p=0.93; axons: 0.38 ± 0.003, F=0.31, p=0.95; global orientation selectivity index, gOSI, boutons: 0.28 ± 0.003, F=0.55, p=0.80; axons: 0.27 ± 0.003, F=0.77, p=0.61; preferred temporal frequency, boutons: 2.70 ± 0.10 Hz, F=1.07, p=0.39; axons: 2.83 ± 0.09 cpd, F=0.53, p=0.81; RF strength, boutons: 1.89 ± 0.02, F=0.77, p=0.61; axons: 1.87 ± 0.02, F=0.61, p=0.74; preferred spatial frequency, boutons: 0.090 ± 0.004 cpd, F=5.87, p=9x10-6; axons: 0.085 ± 0.003 cpd, F=4.17, p=4x10-4. One-way ANOVA.

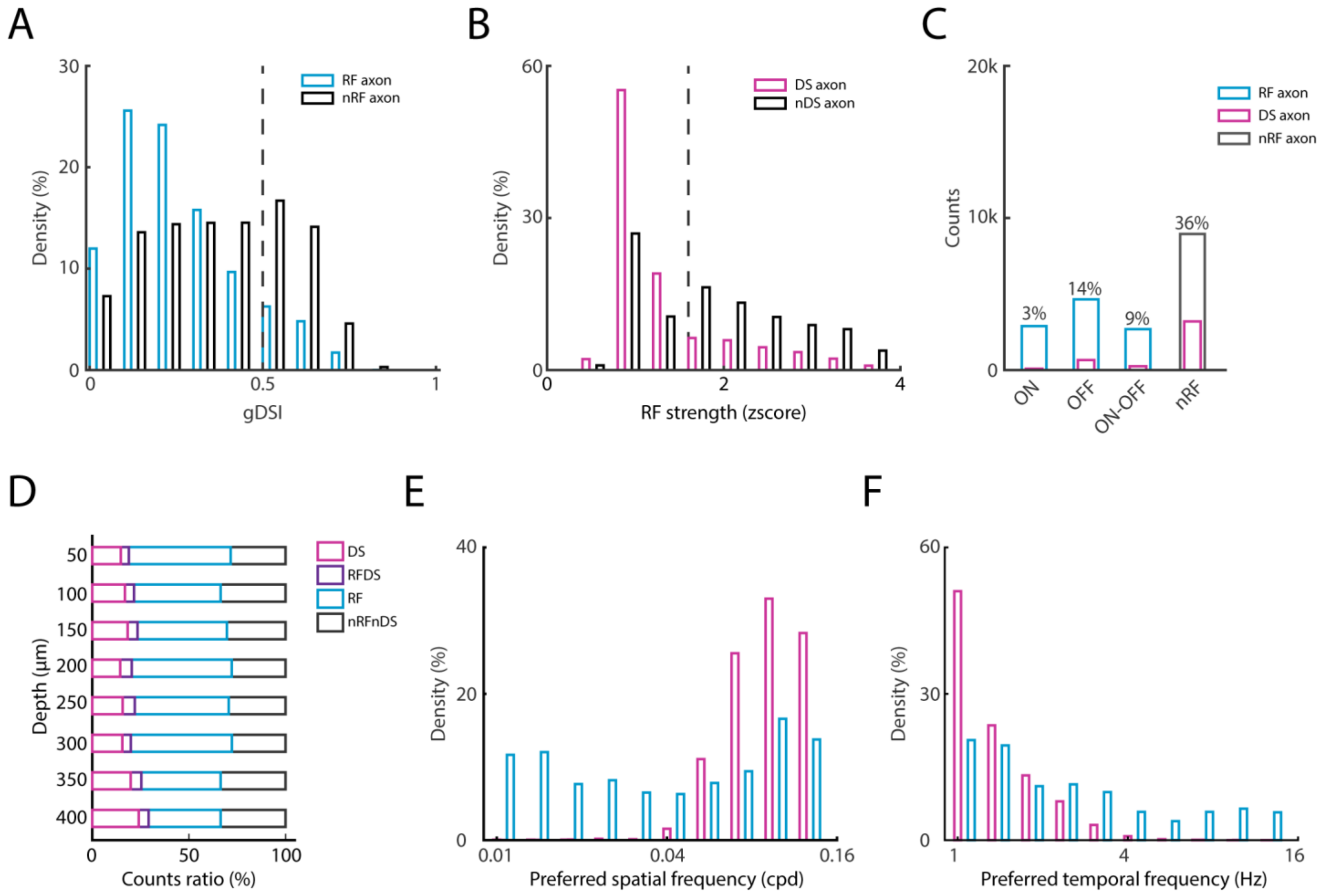


Figure S5. Comparisons between DS and RF axons.

(A) – (C) same as Figure2 (A) – (C) but for axons.

(D) – (F) same as Figure2 (E) - (G) but for axons.

Statisics: DS axons (n=2295) vs RF axons (n=3590), spatial frequency: 0.098 ± 0.035 vs. 0.043 ± 0.061 cpd, U=53118, $p<10^{-100}$; temporal frequency: 1.46 ± 0.59 vs. 2.51 ± 2.70 Hz, U=1917535, $p<10^{-100}$, Mann-Whitney U test.

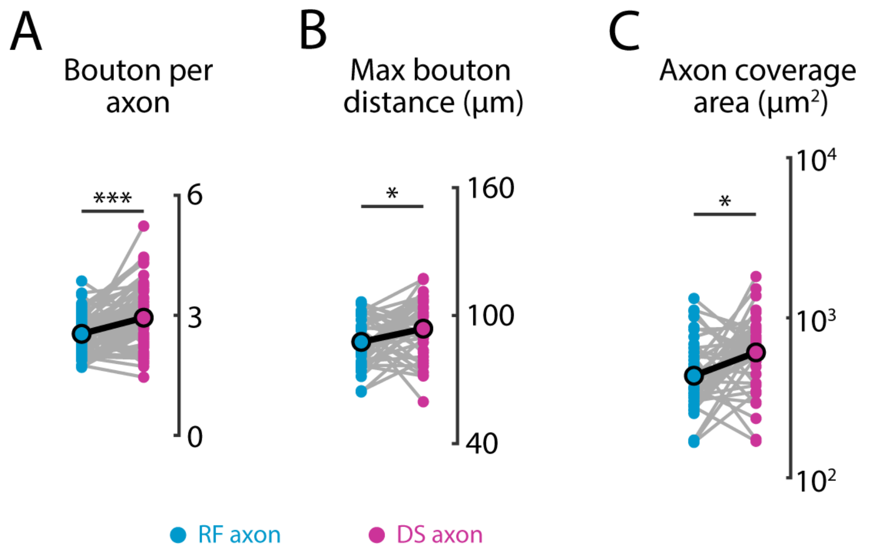


Figure S6. Comparisons of axon morphology estimates between DS and RF axons.

Statistics: DS axon vs. RF axons per plane. Bouton counts: 2.94 ± 0.01 vs. 2.54 ± 0.06, n=70, W=563, p=7.0x10$^{-5}$; max bouton distance: 94.5 ± 2.1 vs. 88.3 ± 1.8 µm, n=41, W=263, p=0.03; axon coverage area: 2362 ± 122 µm$^2$ vs. 2162 ± 107 µm$^2$, n=41, W=236, p=0.01. Wilcoxon rank test.

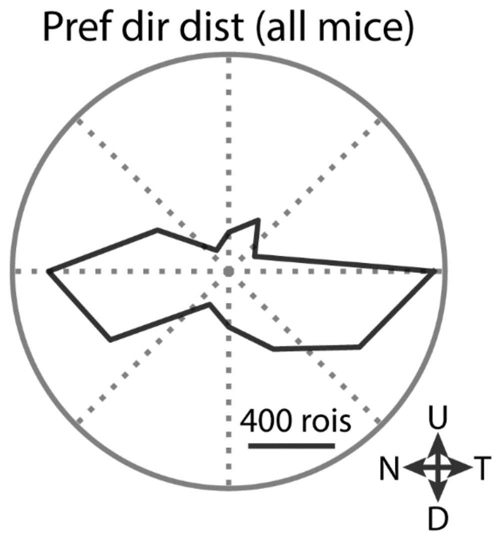


Figure S7. Distribution of preferred directions of all DS boutons.

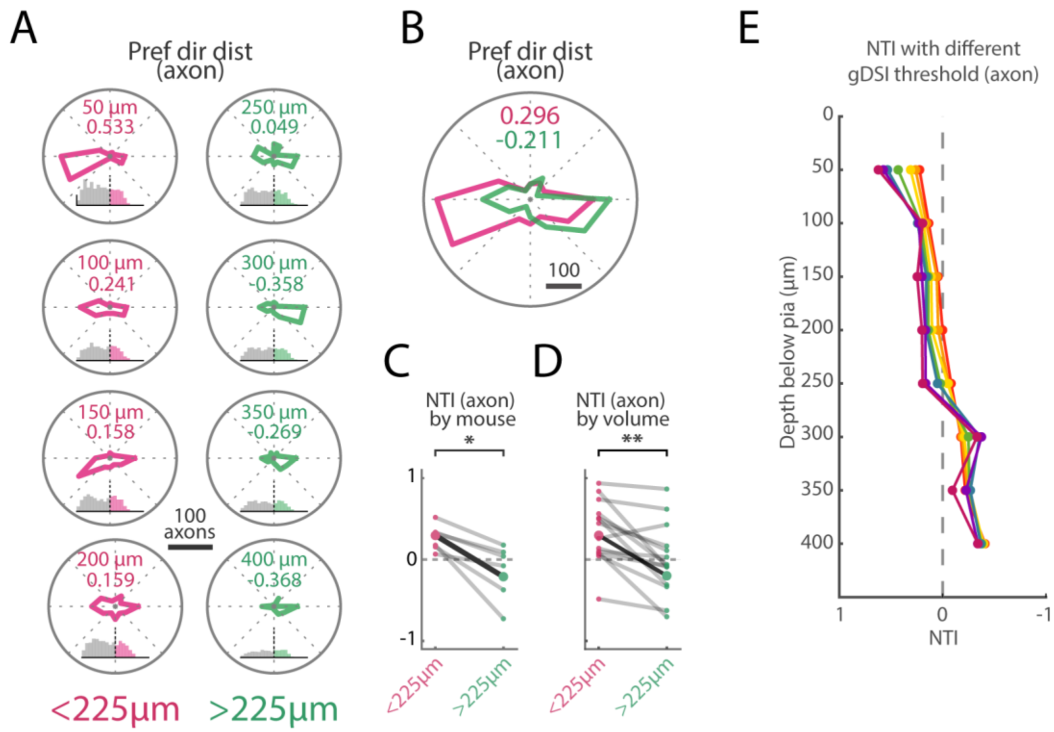


Figure S8. Same as Figure 3 but for axons.

Statisitcs: NTI superficial vs. deep, for mice: n=6, 0.26 ± 0.06 vs. -0.14 ± 0.12, W=0, p=0.028; for volume: n=14, 0.34 ± 0.10 vs. -0.01 ± 0.11, W=6, p=0.0035, Wilcoxon rank test.

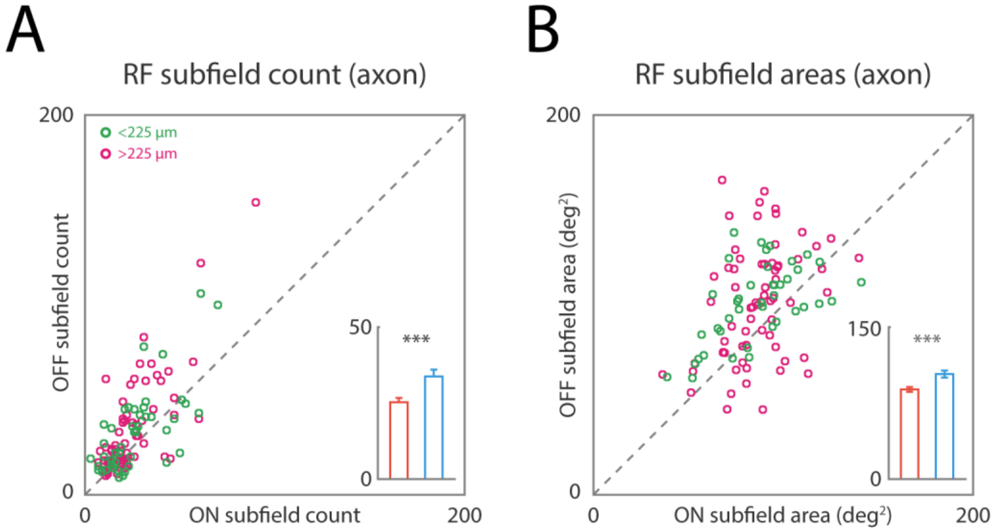


Figure S9. Same as Figure 4 (A) and (B) but for axons.

Statistics: ON vs. OFF, axon counts per plane, all depth: n=116, 25.3 ± 1.41 vs. 33.8 ± 2.23, W=1436.5, $p=1.15x10^{-7}$; superficial: n=70, 24.4 ± 1.76 vs. 33.9 ± 3.05, W=447.5, $p=5.45x10^{-6}$; deep: n=46, 26.7 ± 2.31 vs. 33.5 ± 3.19, W=288, p=0.006. Mean receptive area per plane, all depth: n=116, 85.5 ±1.84 vs. 100.9 ± 2.45, W=1358, $p=2.07x10^{-8}$; superficial: n=70, 85.6 ± 2.25 vs. 100.6 ± 3.48, W=578, p=0.0001; deep: n=46, 85.3 ± 3.14 vs. 101.5 vs. 3.16, W=159, $p=3.07x10^{-5}$. Wilcoxon rank test.

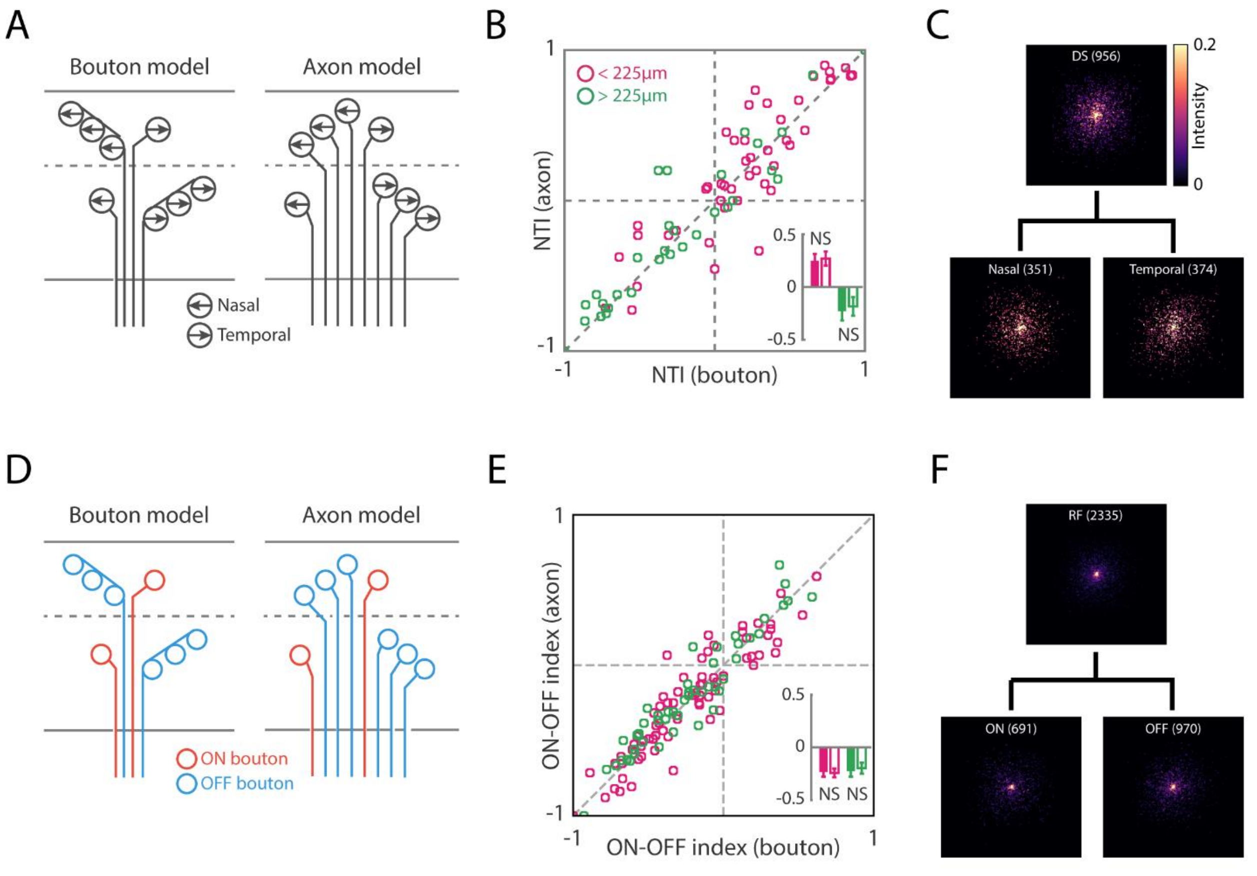


Figure S10. Functional specific bouton distribution can be explained by axon numbers of each group but not bouton number per axon.

(A) Two possible models (bouton model vs. axon model) that can give rise to the depth-dependent bias of motion direction preference.

(B) Bouton vs. axon comparison of NTI for each imaged plane. Inset: bar-graph showing no significant differences in NTIs between boutons and axons at different depths.

(C) Stacked axon masks of nasal preferring and temporal preferring DS axons.

(D) Bouton model vs. axon model for OFF dominance.

(E) Bouton vs. axon comparison of ON-OFF index for each imaged plane. Inset: bar-graph showing no significant difference in ON-OFF indices between boutons and axons at different depths. ON-OFF index: $(n_{ON} + n_{OFF)}$ / $(n_{ON} - n_{OFF})$

(F) Stacked axon masks of ON and OFF RF axons.

Statisitcs: Bouton NTI vs. axon NTI plane-wise: superficial, n=49, 0.25 ± 0.06 vs. 0.27 ± 0.07, W=467, p=0.42; deep: n=31, -0.23 ± 0.09 vs. -0.18 ± 0.09, W=153, p=0.25. Bouton ON-OFF index vs. axon ON-OFF index plane-wise: superficial, -0.235 ± 0.042 vs. -0.243 ± 0.042, n=72, W=1169, p=0.532; deep: -0.225 ± 0.05 vs. -0.198 ± 0.05, n=48, W=376, p=0.072, Wilcoxon rank test.

Video S1. Two-photon images of calcium activities from LGN axons/boutons expressing GCaMP6s in V1 of an awake mouse. Imaging depth: 400 µm. Display speed 10x real time. Field of view: 179.2 x 179.2 µm. The image plane is the same as showed in Figure 1 (E) – (H).
